# Inferring Tumour Proliferative Organisation from Phylogenetic Tree Measures in a Computational Model

**DOI:** 10.1101/334946

**Authors:** Jacob G. Scott, Philip K. Maini, Alexander R. A. Anderson, Alexander G. Fletcher

## Abstract

We use a computational modelling approach to explore whether it is possible to infer a tumour’s cell proliferative hierarchy, under the assumptions of the cancer stem cell hypothesis and neutral evolution. We focus on inferring the symmetric division probability for cancer stem cells in our model, as this is believed to be a key driving parameter of tumour progression and therapeutic response. Given the advent of multi-region sampling, and the opportunities offered by them to understand tumour evolutionary history, we focus on a suite of statistical measures of the phylogenetic trees resulting from the tumour’s evolution in different regions of parameter space and through time. We find strikingly different patterns in these measures for changing symmetric division probability which hinge on the inclusion of spatial constraints. These results give us a starting point to begin stratifying tumours by this biological parameter and also generate a number of actionable clinical and biological hypotheses including changes during therapy, and through tumour evolution.

## Introduction

The cancer stem cell hypothesis (CSCH) posits that tumours are composed of a hierarchy of cells with varying proliferative capacities. Under this hypothesis, a subpopulation of ‘cancer stem cells’, also termed tumour initiating cells (TICs), are able to self-renew through symmetric division and also to differentiate into tumour cells resembling transit amplifying cells (TACs) through asymmetric division (see Fig 1A), giving rise to the entire diversity of cells within a tumour^1^. The CSCH provides a conceptual framework by which to understand many different aspects of cancer progression, including: the occurrence of functional heterogeneity despite genetically identical states^2–4^; resistance to chemotherapy^5,6^ and radiotherapy^7–9^; recurrence^10^; and metastasis^11^. Despite its popularity, the CSCH has been the subject of continual debate and modification in order to maintain compatibility with experimental observations^12–14^.

**Figure 1.**
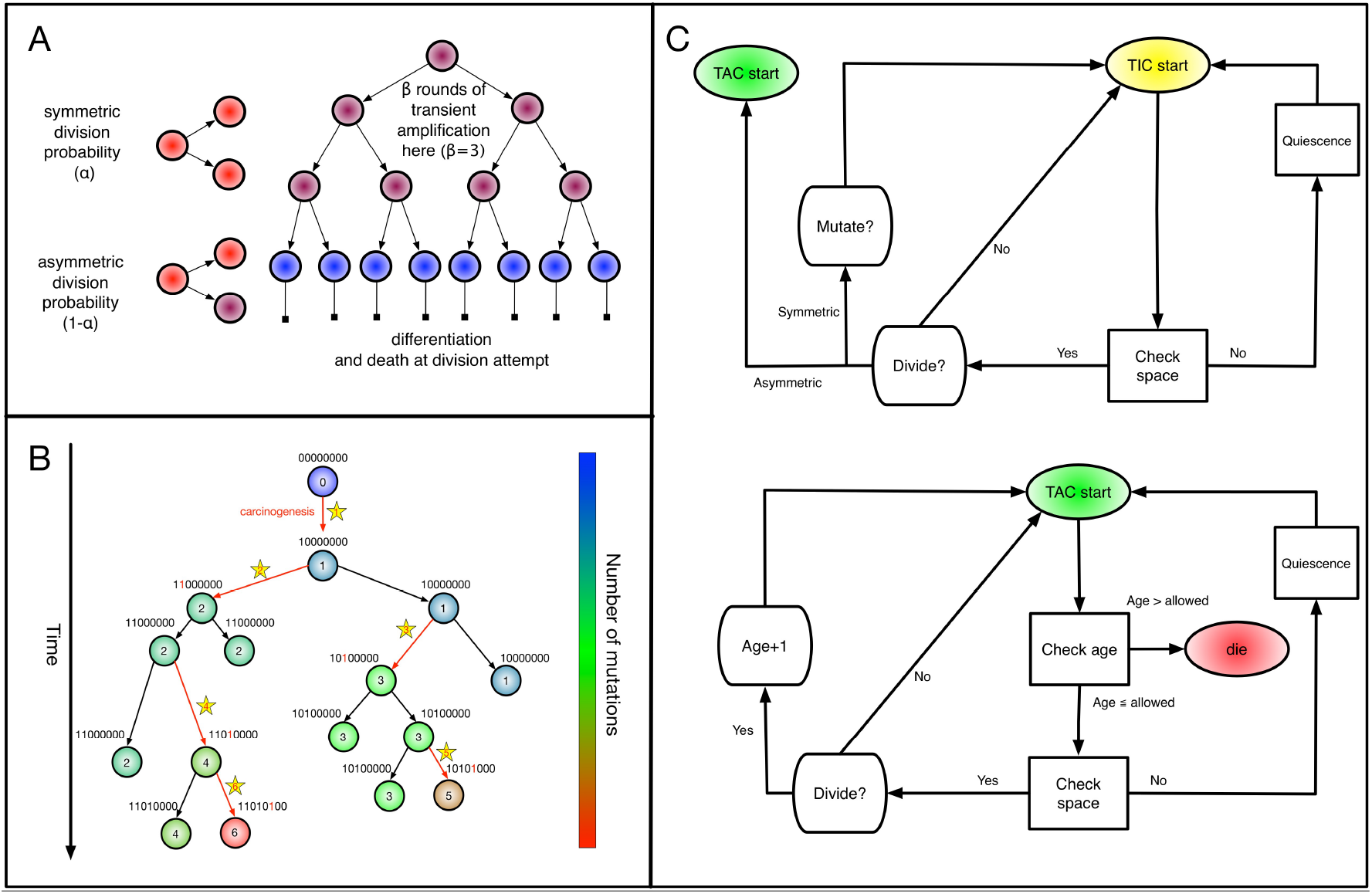
Spatial stochastic model schematic with neutral mutation schema. (A) The proliferative hierarchy. Each TIC can divide symmetrically with probability *α* to make two identical TIC progeny, or asymmetrically with probability (1 — *α*) to make one TIC and one TAC. TACs divide symmetrically until they reach a specific divisional age (*β* = 4 for this work), after which they die upon division attempt. (B) At each division event (branching) after the first (carcinogenesis, labelled with a 1), a random number of mutations drawn from a Poisson distribution with expectation λ is conferred on each daughter (subsequent starred events). Each mutation event is given a unique flag, which is inherited by its offspring unless they too mutate. Each unique mutation can then be considered as a novel mutant allele (red) appearing in the population. (C) Flowchart outlining cellular automaton rules governing TIC and TAC growth, including spatial inhibition of growth and TAC age.

While the specifics of the CSCH are still a matter of debate, the clinical relevance of those cells with traits ascribed to TICs is clear. Regardless of the accepted importance of this knowledge, our ability to measure their dynamics in a clinical setting is lacking. In *vivo* measurement efforts are limited to carefully conducted live imaging in genetically engineered mice^15^, or genetic labelling and subsequent lineage tracing^16^; while in *vitro* systems are better suited to the extraction of these parameters, little has been done to quantify them, as technically demanding single-cell lineage tracing^17^ is required. These experimental difficulties speak to the need for more theoretical work in this area, especially to propose metrics for quantifying proliferative parameters such as TIC symmetric division probability (Fig 1A) from clinical data. This is of particular importance as there is mounting evidence for the relevance of a proliferative hierarchy in determining response to radiotherapy^18^ and chemotherapy^5^. Further, we now know that certain microenvironmental factors such as hypoxia^19,20^, acidosis^21^, growth factors^22^, and even stromal cell co-operation or co-option^23,24^, can perturb this system.

Several published mathematical models, taking different forms and considering different aspects of heterogeneity, have predicted that the evolution of a solid tumour should depend strongly on whether or not it exhibits a proliferative hierarchy, and on the parameters of such a hierarchy. These models have included spatial proliferation constraints, microenvironmental heterogeneity and selective pressures, and the noted differences include shape, clonal heterogeneity, rate of evolution and growth dynamics. Werner at al. specifically studied the differences in bulk tumour behaviour between tumours arising from mutant TICs and TACs^25^ in a non-spatial context. In a spatial context, the work of Sottoriva et al.^3,26^ and Enderling et al.^27,28^ represent the first works in which it was shown that the parameters governing TAC dynamics can constrain tumour growth, and also to show that TIC-driven tumours have significantly different spatial growth patterns: specifically, that they exhibit ‘patchy’ growth. In none of these models, except Sproufsske et al.^29^, in which the main question centred on TAC numbers, were these differences studied across TIC symmetric division probabilities, which is a key parameter governing the hierarchy, and one that is exceedingly difficult to measure or perturb in *vitro* or in *vivo.*

To describe the evolutionary relationship between members of a species, or larger groups of life forms, biologists often formulate tree diagrams that represent their specific hypotheses about relatedness. While tree diagrams have been in use since medieval times to describe genealogies, their use to describe animal species was not popularized until the early 1800s. These trees were originally made on the basis of gross morphological differences (or similarities) and were called phenograms or cladograms, but in the last few decades we have begun to define these differences based on genetic information. The field of phylogenetics, born in the 1980s, seeks to use objective, genetic information to build trees. When populations are sampled, a common method of understanding the clonal evolution is through phylogenetic reconstruction, a method of inferring, usually from genetic sequence similarity, the evolutionary life history of a given life form. This has classically been applied in scientific fields such as zoology, and it has become a branch of bioinformatics all of its own, even spawning a branch of discrete mathematics called T-theory^30^.

Phylogenetics has, in the last decade, begun to be applied to cancers, giving rise to a subfield recently dubbed ‘PhyloOncology’ by Somarelli and colleagues^31^. Using phylogenies reconstructed from spatially separated biopsies and informatic algorithms, many aspects of tumour evolution have begun to be elucidated^32^, including the genetic heterogeneity present within a primary tumour^33^, the origin of individual metastatic tumours within the primary site^34,35^, and the effect of chemotherapy on primary and metastatic sites^36,37^.

In addition to these sorts of questions, there are precedents in other fields for using phylogenetic information, integrated with population dynamics, a technique called phylodynamics^38^, to infer other underlying biological processes. For example, Leventhal et al.^39^ proposed that the phylogenetic tree contains a ‘fingerprint’ that can be used to determine the evolutionary process driving the population in question. Modelling the spread of HIV within a contact network, the authors investigated whether the network structure could be inferred from the resulting disease phylogenies. To address this question, the authors simulated a range of epidemics on several families of random graphs and measured the resulting phylogenetic trees, finding that certain tree-based measures could discriminate between the qualitatively different families of random graph structures considered.

We hypothesize that a similar approach could be used to discriminate between in *silico* tumours with different symmetric division rates. To test this hypothesis, here we study the effect of TIC symmetric division probability on tumour evolution using a computational modelling approach. We focus on observed patterns in reconstructed phylogenetic trees across a range of symmetric division probabilities. The estimation of this proliferative parameter from clinical data could help improve our understanding of the effect of therapies on tumour growth dynamics, and our ability to stratify tumours for consideration of different therapies. In this way, we seek to provide translatable measures to aid in understanding tumour biology: to use mathematical modelling to ‘see the invisible’.

The remainder of this paper is structured as follows. We first present a spatial stochastic model of tumour growth under a proliferative hierarchy with neutral mutations, which we embed on a twodimensional lattice to enable the study of the effect of spatial constraints. Next, we develop an algorithm to reconstruct the branched phylogenetic structure from each realization of our tumour growth model. We apply a range of statistical measures of phylogenetic tree shape to simulation outputs for comparison. We explore the temporal dynamics of these measures over the course of tumour growth to assess whether they are robust to tumour size changes, and then to changes in mutation frequency. Finally, we discuss the possible clinical utility of these measures.

## Materials and Methods

### Model development

Here, we describe the development of a two-dimensional, lattice-embedded cellular automaton (CA) model of tumour growth with contact inhibition growing under neutral evolution and a proliferative hierarchy.

#### Proliferative hierarchy

We model a proliferative hierarchy comprising two cell types, TICs and TACs. We assume that each TIC divides symmetrically with probability *α*, creating two TICs, and asymmetrically with probability 1 – *α*, creating one TIC and one TAC. While there is evidence that microenvironmental parameters such as nutrient deprivation^40^, acidity^21^ and hypoxia^41,42^ can change symmetric division probability, and that it is likely to vary from cell to cell, for simplicity we will assume it is constant. As it has been shown theoretically that the overall population dynamics of TIC-driven tumours is equivalent with or without TIC symmetric differentiation^43^ (when a TIC divides to create two TACs), and as the lineage extinction possible in this case would significantly complicate our phylogenetic analysis, we make the simplifying assumption that there is no symmetric differentiation. We do not rule out that the addition of symmetric differentiation could affect phylodynamics, but leave that question for further study.

We assume that every TAC division is symmetric, creating two TACs, but only allow this to progress for *β* rounds of division, after which the TAC will die if chosen to divide again. Here *β* represents the replicative potential of TACs, and is posited to represent telomere length^44^. Previous theoretical work has shown that tumour growth kinetics in spatially constrained geometries are strongly affected by the value of *β*^28^. In particular, if *β* > 5, then simulated tumours experience unrealistically lengthy growth delays. Therefore we follow a previously used assumption^3,29^ and fix *β* = 4. This mode of growth and differentiation is illustrated in Fig 1A. For simplicity, we neglect cell death, though this could be added as a straightforward extension in future work.

#### Neutral evolution

To understand the effects of neutral evolution on tumours with differing proliferative hierarchies, we extend our model of tumour growth under a proliferative hierarchy to include random mutations. At each cell division, there is a possibility that one or more mutations occur. To determine the number of mutations accumulated by a given daughter cell, we independently draw a random number from a Poisson distribution with rate λ. We assume for simplicity that every mutation arising in our model is unique. This ‘infinite sites’ assumption is usually ascribed to Kimura^45^.

For simplicity, we assume that mutations confer no advantage, disadvantage or any other phenotypic change and therefore serve only as a method by which to track clonal lineages. This assumption could in principle be loosened to allow for positive selection^46^, a balance of positive and negative selection^47^, and neutral evolution^48^. A schematic of this model of evolution, and labelling scheme, is shown in Fig 1B.

For computational efficiency, we record a unique flag only for the most recent mutation accumulated within a cell, which is passed down to its progeny, unless a mutation occurs, in which case a new flag is assigned. We also record each mutation event in the form of an ordered pair (parent flag, child flag), so that the complete ‘genomes’ (bit strings) can be reconstructed for future use. As they are the only cells capable of forming tumours on their own, and infinite replication, we follow previous works in considering new mutations to accrue only in TICs^3,26,29,49^.

#### Spatial dynamics

As we are interested in the effect of the proliferative hierarchy on the neutral evolutionary process in solid, spatially constrained tumours, we embed our cell-based model in a two-dimensional square lattice. While recent work has shown some qualitative differences in vascularised CA models between two and three dimensions, using a two-dimensional lattice for unvascularised tissue is a common simplification^50–53^ that allows spatial constraints to be studied in a computationally tractable manner. In addition to the above description of cell proliferation, we consider cell proliferation to be modulated by contact inhibition^54^. Each cell is allowed to divide only if there is one or more free lattice sites within that cell’s Moore neighbourhood; if not, then we consider the cell to be in a quiescent state that may be exited when space becomes available. At each time step, each ‘cell’ has an opportunity to divide given that it has space to do so. Cells are chosen uniformly at random for updates from the entire population to avoid order bias.

#### Cell-type specific rules

If space is available, and the cell is a TIC, then the type of division is determined by choosing a uniform random number, r, from [0,1]. If *r* < *α*, then the TIC divides symmetrically, creating another TIC that is placed uniformly at random in one of the free neighbouring lattice sites. The parent and daughter TICs will independently acquire a random number of new mutations, as described above. If *r* ≥ *α*, then the TIC divides asymmetrically, creating a TAC that is placed uniformly at random in one of the free neighbouring lattice sites. The daughter TAC is created with the same mutation ID as the parent, and age = 0, while the parent TIC will independently acquire a random number of new mutations, as described above.

If the chosen cell is instead a TAC, then the check after available space is a check of the cell’s proliferative age, which is the number of divisions as a TAC. If the TAC age is equal to the replicative potential, *β*, then the TAC dies, at which point it is removed from the simulation. If the TAC age is less than *β*, then we create a new TAC daughter and place it in an empty space in the Moore neighbourhood at random. The parent and daughter TACs share the same mutation ID and their age is updated to be one more than the age of the originally chosen TAC.

#### Full implementation

The full CA flow-chart, represented in Fig 1C, schematises the entire process of cell fate decisions that each cell undergoes at each time step in the spatial model. In the top panel, the rule set followed by the TICs is represented to include differentiation and mutation. In the bottom panel, the TAC rule set is defined to include death by terminal differentiation and TAC aging. An example simulation of tumor growth over time is shown in Fig 2, where the effect of lowering *α* can be seen on overall tumour growth kinetics, where the colour-bar represents the current clonal state (mutation ID) of a given clone.

**Figure 2.**
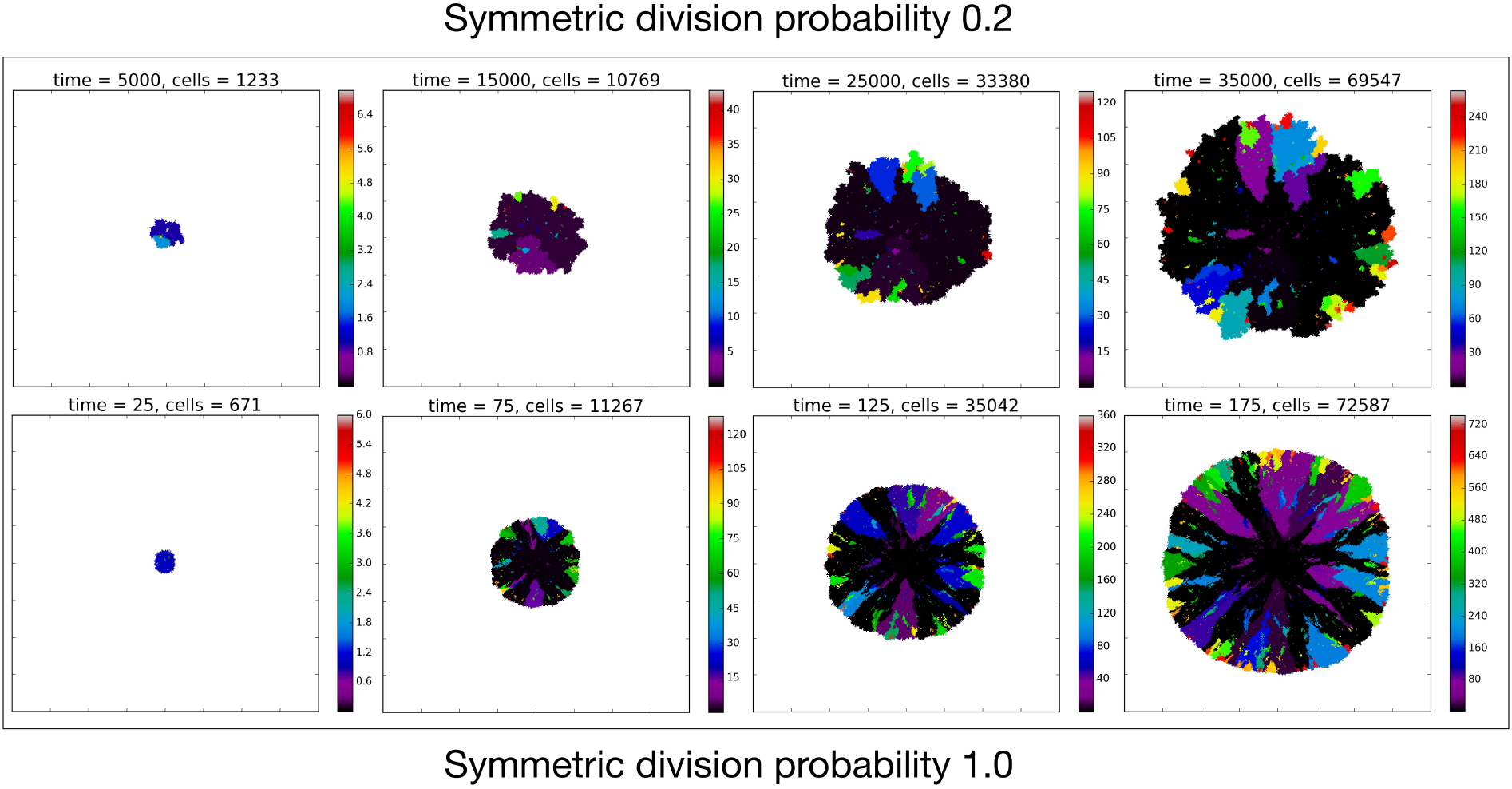
Temporal evolution of the spatial model reveals observable morphologic differences between TIC-driven and non-TIC-driven tumours, as observed by others. We plot representative results of simulations of two tumours, each simulated on a square lattice of size 400 × 400. Top: a tumour simulated with *α* = 0.2 and *β* = 4. We notice, as have Enderling et al.^27^ and Sottoriva et al.^3^, a ‘patchy’ clonal architecture, and non-uniform edge. Bottom: a tumour simulated with *α* = 1.0, i.e. no proliferative hierarchy. We note smooth edges, radial patterns of clonal architecture and relatively faster population growth, reaching ≈ 70,000 cells in less than 200 time steps. To reach a similar size, the tumour with symmetric division probability of 0.2 took 35,000 time steps. Colour bars denote number of mutations present in a given clone, note that the top scale is about 1/3 of bottom scale.

### Recovering phylogenetic trees from simulation

While experimentalists and clinicians can only infer phylogenies from incomplete data, reconstruction of the ‘true’ phylogeny is possible in our model as we can record the entire life history of the simulated tumour. Thus, we can test whether phylogenetic tree-based measures are able to discriminate TIC symmetric division probability in the case where the ‘ground truth’ is known. At each time step we record the spatial location of each individual cell with its mutation ID, which is our CA state vector. Additionally, we record the evolutionary ‘life history’ as a list of ordered pairs of every mutation event (parent mutational ID, child mutational ID). We then recursively construct the phylogenetic tree from this life history.

#### Phylogenetic tree reconstruction algorithm

To create the complete tree data structure required for our quantitative analyses we use the information encoding the mutation events from our stochastic simulation. To this end, we create a list of unique parent-child pairs using the life history of mutation events. We then apply an iterative process in which each child is added as a subnode below the parent (from the unique parent-child pair). This process is continued until all parent-child pairs are added to the structure, and the tree is complete. The simulation code and functions to create these trees and calculate the metrics is freely available on request.

#### Qualitative comparison of reconstructed trees

To compare phylogenies from simulations with different underlying parameter values, we first construct and visualize the phylogenies constructed from three example simulations with differing TIC symmetric division probabilities in Fig 3. It is clear by inspection that the number of mutations increases with symmetric division probability (more branches). However, the tree structure is not as easy to parse visually. For ease of visualization the trees depicted in Fig 3 have been pruned of all terminal nodes (also called leaves) with no children of their own. While this transformation does affect the quantitative results, it does not qualitatively affect the resultant phylogenetic tree statistic ranks (see Fig 8). All analyses shown will utilize the full trees.

**Figure 3.**
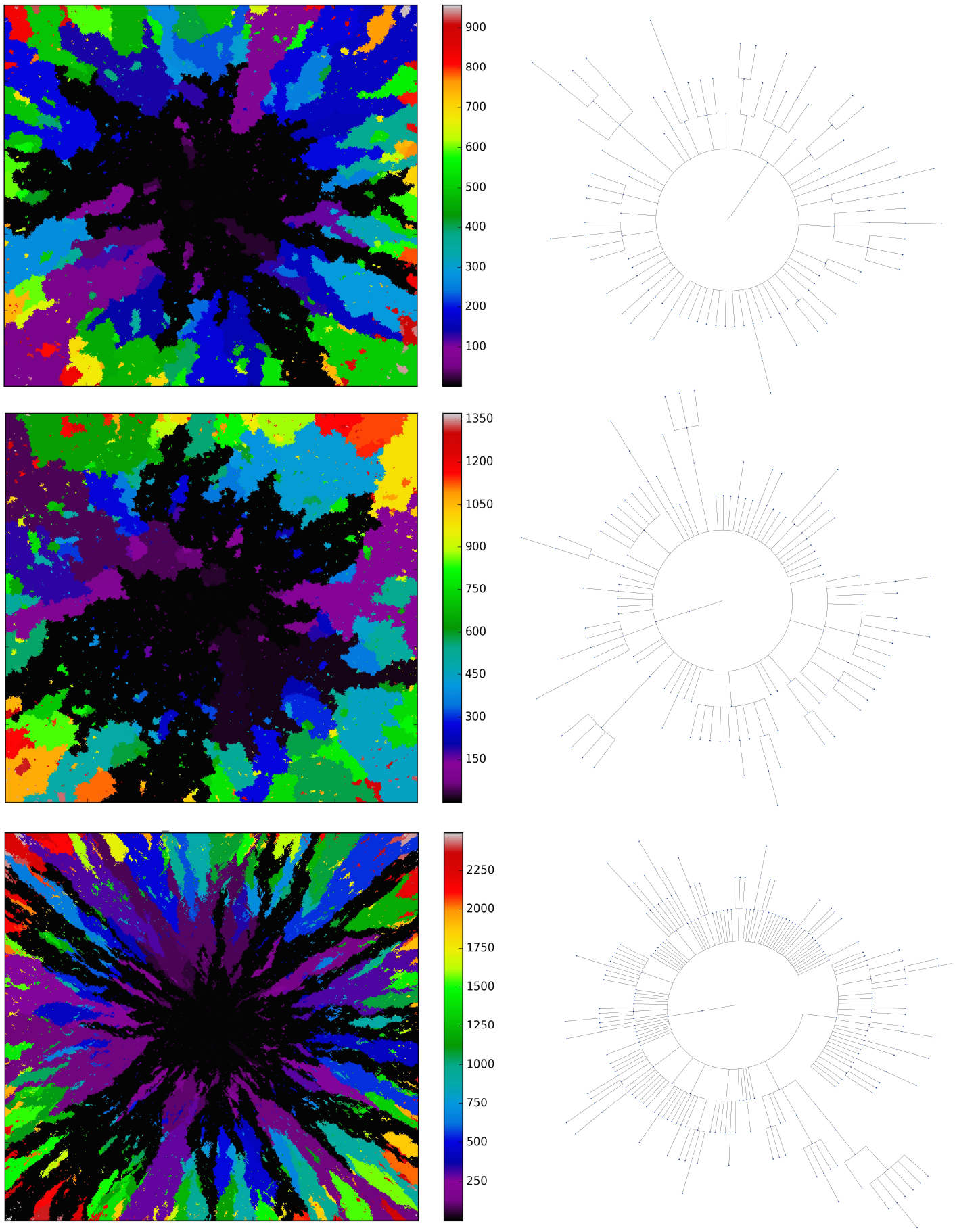
Three example simulations with increasing symmetric division probability, *α* (0.2, 0.6 and 1.0 from top to bottom) and their associated phylogenetic trees. Each example plot is the result of a single stochastic simulation of our spatial CA model. Each simulation is initiated with a single TIC and complete when the domain is full, in this case 250,000 cells. Parameter values are *β* = 4 and λ = 0.01. Visualized trees (right) have been pruned of all leaves for ease of visualisation, which does not qualitatively affect measure rank (see Fig 8).

### Candidate tree-based measures for model comparison

Visual inspection of Fig 3 suggests that simulations with different TIC symmetric division probabilities generate distinct phylogenetic trees. However, to make meaningful conclusions we must perform a quantitative comparison. Here we present several measures useful in summarising and comparing phylogenetic trees. The most commonly studied property of a phylogenetic tree’s shape is its balance, defined as the degree to which internal nodes (branch points) have the same number of children as one another. Balance (or imbalance) indices depend only on the branching topology of trees, and not on other factors like branch length or other features of the terminal branches (leaves). Since the first balance index by Sackin^55^, many others have been proposed with slightly differing properties^56^. One of the first papers to present a systematic comparison of a suite of balance indices (often denoted with the letter ‘B’) and indices of imbalance (denoted with ‘I’) was by Shao and Sokal^57^, who reported striking differences between the studies’ measures. Their central message was that different measures on trees can give insight into different aspects of the underlying processes governing the interactions, and one should thus consider several measures for any given tree or family of trees. In this study we will consider several tree topology-based measures.

Before describing the measures, it is worthwhile to briefly define the terms which are used to describe trees, and the two basic underlying stochastic models which have been proposed to describe neutral evolution and the resulting topologies. Phylogenetic trees are mathematical objects which describe the evolutionary relationship between individuals with different physical traits from one another, or in the case of our model, different mutational combinations (genotypes). In our model, each simulation begins with a cell with mutation flag 1, or a genotype with the first allele mutated (1000…), termed the ‘root’, and evolution progresses stochastically, by adding individual mutations at subsequent alleles and increasing the mutation flag, as described in Fig 1B. At each mutation event, an evolutionary branch point is created, which is termed a node in phylogenetic tree terminology. If this node gives rise to no other children during the simulation, it is termed a terminal node, or leaf. There are two common, classically referenced models, which bear mention here as well, since many tree topology-measures are normalized against them. The first, described by Yule in 1924^58^ and sometimes termed the ‘equal rate Markov’ model, begins with a single root and proceeds by replacing, uniformly at random, a given leaf with a node with two children of its own. The process continues until the desired number of leaves exist. The other main model, termed the ‘Proportional to Distinguishable Arrangements’ or uniform model, was described by Rosen^59^. This model, which is truly a model of tree growth rather than an explicitly evolutionary process, begins as does the Yule model (and indeed ours) with a single node labelled 1. At each update step, a new leaf is added to the tree at any point, either internal node or leaf. These models will serve as normalisation factors in several of the measures we present below, which are summarised graphically in Fig 4.

**Figure 4.**
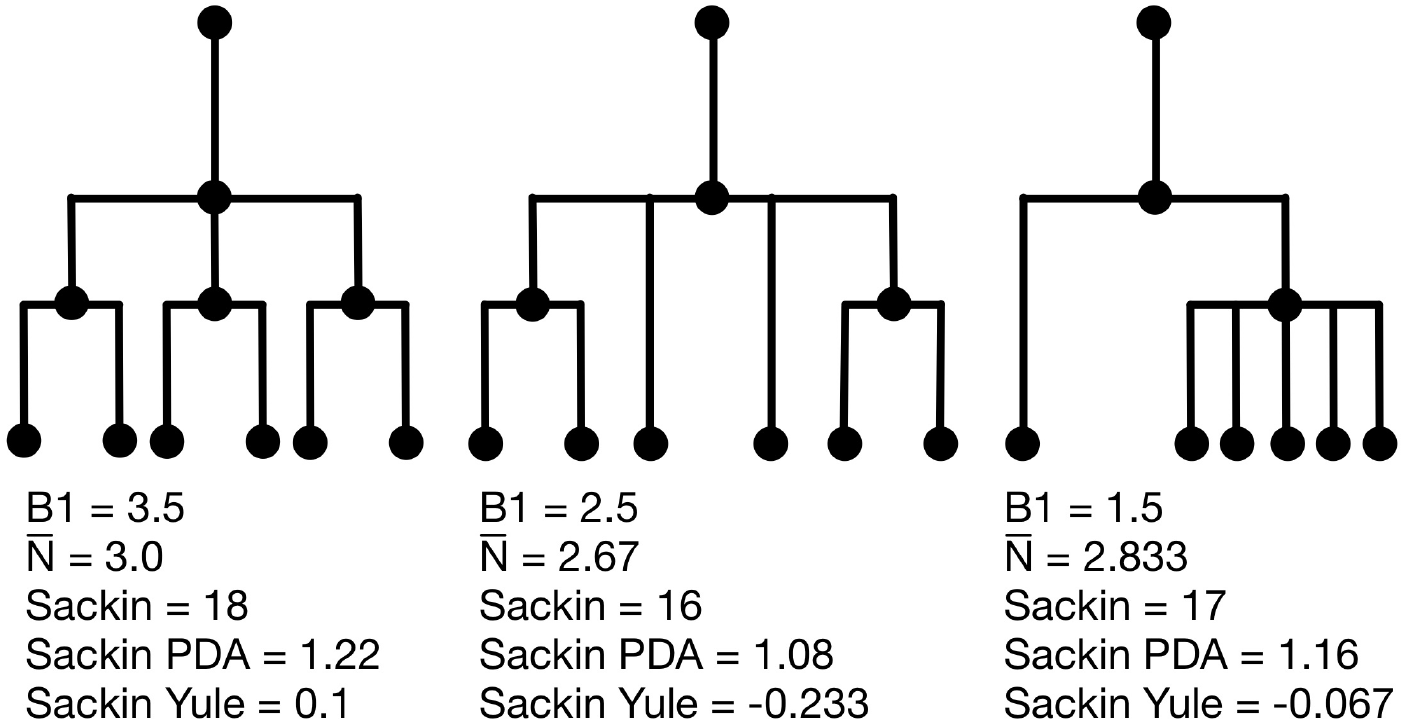
Example phylogenetic trees and their measures. From left to right the trees contain 4, 3 and 2 internal nodes (dots) respectively, but the same number (6) of terminal nodes.

#### Sackin index

The Sackin index was the first statistic used to understand the balance of a phylogenetic tree^55,57^. To compute this statistic, one sums the number of ancestors (*N_i_*) for each of the n terminal nodes of the tree:

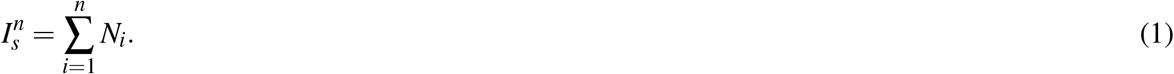

This index increases with tree size: under the Yule growth model, its expectation 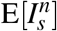 grows as 2nlogn^58^. One can therefore only perform a meaningful comparison of Sackin indices of trees generated from tumours if they are the same size.

#### Normalized Sackin index

To address this dependence on tree size, several normalisations to the Sackin index have been proposed, two of which we explore here. In particular, one can normalise the Sackin index of a phylogenetic tree to the expectation value of a similarly sized tree, under the Yule growth model:

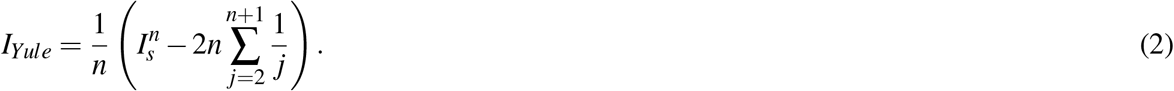

One can alternatively normalise using the Proportional to Distinguishable Arrangements (PDA) model^59,61^ which is simply the Sackin index scaled by *n*^3/2^.

#### The B1 statistic

The B1 statistic, originally described by Shao and Sokal^57^, considers the balance of a tree. To calculate the measure, one uses all *i* internal nodes of the tree with the exception of the root (the founding cell). For each non-root internal node *j*, the maximum number of nodes traversed along the longest possible path to a terminal node, *M_j_*, is counted. The B1 statistic is then defined as

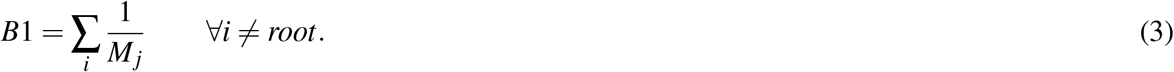

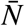 reports the average number of nodes above a terminal node. To compute this, we sum the path from each terminal node to the root, and divide by the number of terminal nodes. An alternative definition is the Sackin index ‘normalised’ by the number of terminal nodes. For a more complete review and comparison of the measures presented here, and others, see Blum et al.^62^ and Shao and Sokal^57^.

Examples of how these measures change on several example trees with equal numbers of leaves (but different numbers of internal nodes) are presented in Fig 4. In these examples, we compute each of the presented measures for comparison. From left to right, the trees contain 4, 3 and 2 internal nodes respectively, but the same number (6) of leaves. We note that the measures do not all follow the same pattern. For an exhaustive description of all possible trees with 6 leaves, and the correlation of a larger family of associated measures, see Shao and Sokal^57^.

## Results

### Measuring trees from simulation

As our primary goal is to identify whether tree-based measures allow discrimination of simulated tumours with different TIC symmetric division probabilities, we focus on changes in tree measures as we vary comparable simulations changing only this parameter. To compare the model tree measures, we first perform 50 stochastic simulations of our spatial CA using a range of TIC symmetric division probabilities (0.2,0.4,0.6,0.8 and 1.0), holding mutation rate and TAC lifetime constant (*λ* = 0.01 and *β* = 4). For each simulation, we construct the resulting phylogenetic tree at tumour size 250,000 cells, as described in the Materials and Methods section. We then measure the value of each summary index defined earlier for all 50 simulations at the final time point and plot the distribution in a box-whisker plot, which is shown in Fig 5 with each data point overlaid in a swarm. Differences between distributions were determined using the Wilcoxon rank sum test. While these statistics were performed post hoc, we should note that standard statistics can be misleading for simulation based studies with arbitrarily large sample sizes^63^ (see Supplementary Fig 9 for effect size).

**Figure 5.**
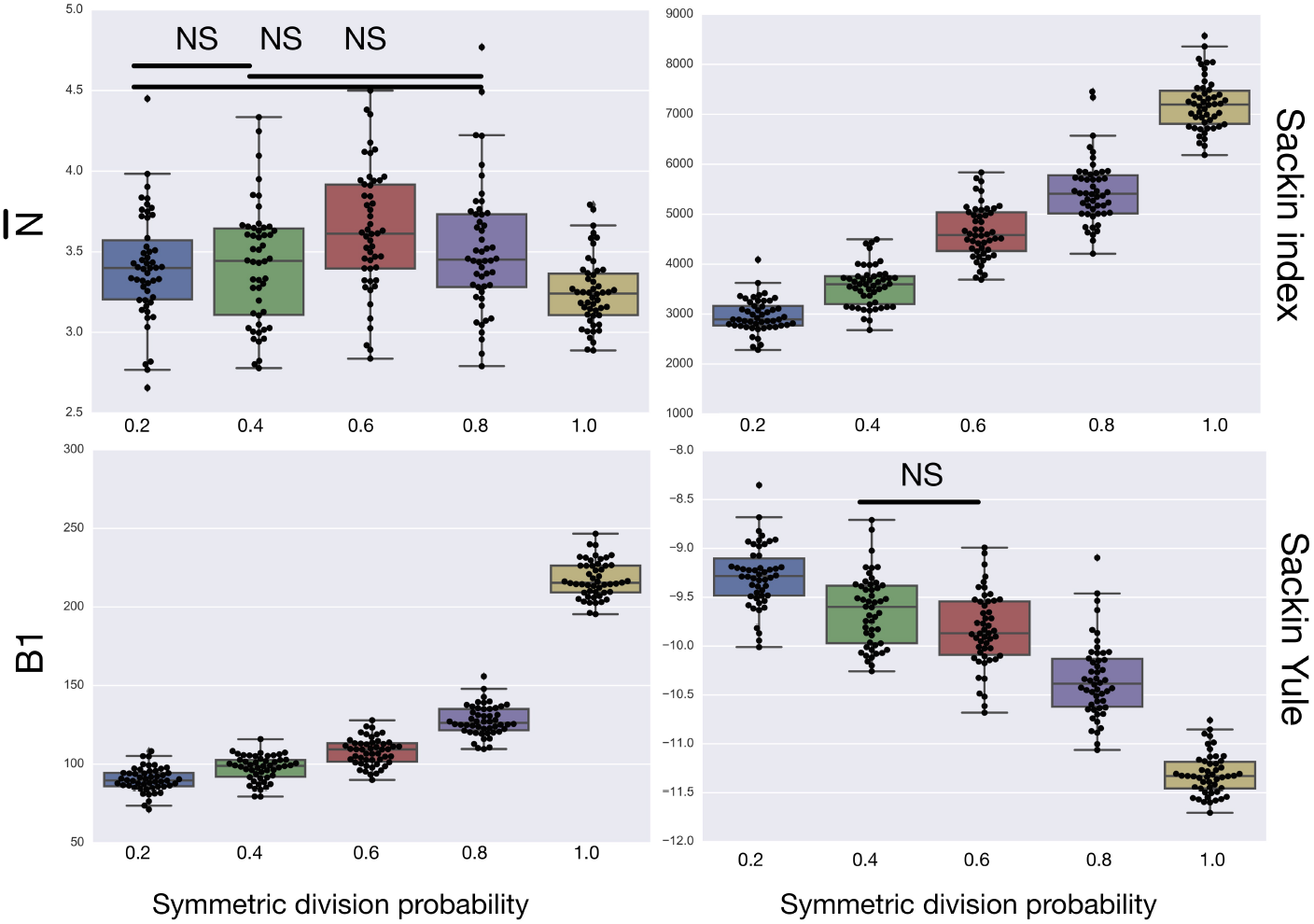
A summary of four tree indices measured over a range of symmetric division probability. We plot the distribution of each of four measures of tree balance for the final resultant trees from 50 simulations against symmetric division probability. All simulations were run with *β* = 4 and λ = 0.01 until a tumour size of 250,000 cells. In each plot we display a box-whisker plot as well as the individual results as points. NS = non-significant by the Wilcoxon rank sum test.

### Variation of tree-based measures with symmetric division probability

The results of the model are presented in Fig 5. We find that all of the indices have monotone relationships with symmetric division probabilities except for 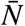 Of the normalised indices, the B1 statistic has the least overlap in error between symmetric division probabilities. All measure distributions are significantly different by the Wilcoxon rank sum test (*p* < 0.05) except 0.4 and 0.6 in the Sackin index normalised by the Yule model (*p* = 0.08). While we recognize the dangers in reporting p-values in simulation based studies^63^, we report them here for comparison, and report effect size as well, with full statistics reported in Figure 9. The strongest effect is seen in the Sackin index (*R*^2^ = 0.871), followed closely by the Yule normalised Sackin index (*R*^2^ = 0.743).

### Dynamics of tree-based measures during tumour growth

As discussed in Materials and Methods, the measures considered here are strongly dependent on the total number of nodes in the tree. With all other parameters held constant, simply allowing a tumour to grow larger would increase the number of total mutations, and therefore the number of total nodes, subsequently altering the value of the measure. To ensure that the differences we have noted are robust to changing tumour size, we next consider how these measures evolve during the growth of a tumour.

To determine how these measures vary over the life of a growing tumour, we measure the index over the course of each simulation at increasing tumour sizes. To accomplish this, we use the life history to reconstruct the tree at 20 equally spaced time points during the lifespan of each of the 50 simulations for each symmetric division probability. The time to fill the domain for each of the symmetric division probabilities is quite different as the dynamics of tumours driven by differing symmetric division probabilities are different (see Fig 2). So, we break the life history into equally spaced time intervals, as the total times in each family of simulations are different. When we compare across symmetric division probabilities we need to consider this ‘time’ to be a surrogate for tumour size instead instead of explicitly comparing times. Comparing across tumour size is of greater utility clinically, however, as the age of a given tumour is rarely known, while size can be readily approximated.

After reconstruction, we then create a ‘time’ trace for each statistic. We plot these statistics over ‘time’ in Fig 6, where each family of 50 simulations (for a given symmetric division probability) is represented by a single trace with the standard deviation represented by the coloured error bars. We find that for each of the statistics, except 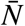, the relationships between the symmetric division probabilities are maintained over time, suggesting that, if we know the tumour size, and true phylogeny, we can estimate the relative symmetric division probability between two samples from these measures. This statement must be somewhat qualified by the fact that mutation probability was also held constant for these simulations. While estimating mutation probability is not trivial, significant advances have been made in measuring the speed of the ‘evolutionary clock’ of tumours: essentially a proxy for mutation probability^64^. Further, we found that the rank order of each discriminatory measure holds throughout tumour growth, indeed becoming more discriminatory as the tumours grow larger (with the exception of 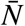). As the tumours simulated in this study are unrealistically small given the computational constraints, this information gives us hope that in tumours of realistic size, these measures would be even more useful. This becomes particularly important as the statistics that we have calculated come from the ‘true trees’, that is, trees comprised of all mutation events. In reality, trees would be inferred from the imperfect information gleaned from biopsies.

**Figure 6.**
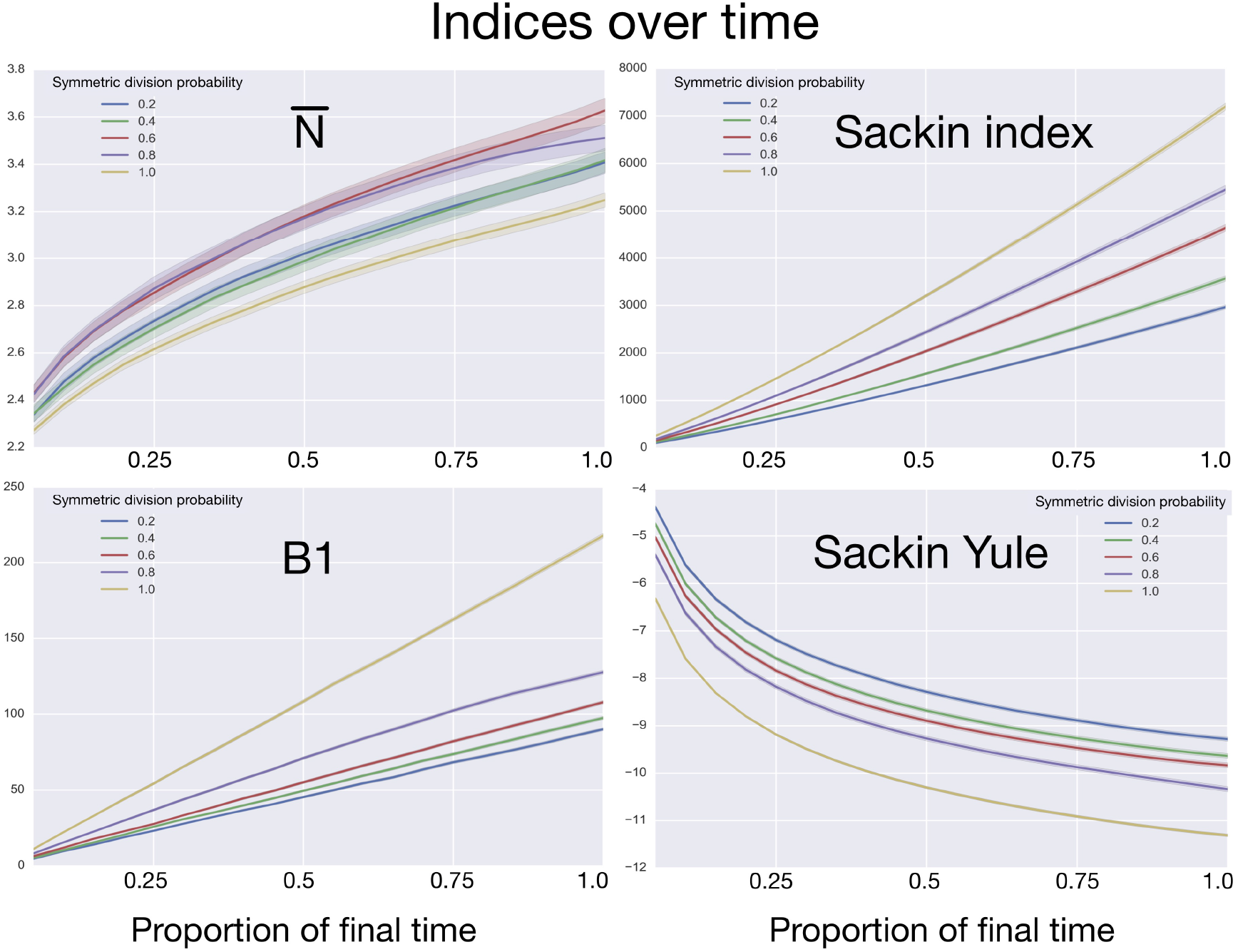
Comparing phylogenetic tree measures across symmetric division probability through tumour growth. We plot the average and standard deviation (error bars) of four phylogenetic tree measures for each of the 50 simulations for a range of symmetric division probabilities over the course of tumour growth. Rank is maintained across symmetric division probabilities for each of the 3 tree measures with which we could discriminate between symmetric division probabilities. As before, 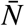 is not predictive and changes rank throughout tumour growth. All tumours are grown to eventual confluence at 250,000 cells. In all simulations *β* = 4 and λ = 0.01.

### Dependence of tree-based measures on mutation probability

As the tree measures depend heavily on the number of mutations within a given tumour, and therefore the number of branches within a given tree, we next ask how these measures behave when we vary mutation probability (λ) and symmetric division probability simultaneously. To answer this, we perform 10 stochastic simulations for each combination of the symmetric division probabilites considered previously and 5 different values for λ varying over two orders of magnitude (0.001,0.005,0.01,0.05,0.1). We then use the previously described method to reconstruct the resulting phylogenies and calculate the measures previously discussed. In particular, we ask how the Sackin index, the B1 statistic and the normalized Sackin index perform over this range of λ to better understand the applicability of these measures in determining differences in symmetric division probability.

We plot the results of this parameter investigation in Fig 7. In each heat map, we plot the mean of the 10 simulations for each parameter combination with symmetric division probability varied along the horizontal axis and mutation probability along the vertical. The indices which are not normalized by branch number, namely the Sackin index and B1 statistic, increase monotonically with mutation probability and symmetric division probability in all cases. The Sackin index normalised by the PDA model, however, varies somewhat unexpectedly and has a global minimum at symmetric division probability of 1.0 and mutation probability 0.01. This measure is monotonic in symmetric division probability except at the highest mutation probability where it becomes somewhat more difficult to determine the differences. As before, the B1 statistic appears to be the most stable, and only breaks down slightly in its ability to distinguish between the families of simulations at the lowest mutation probability (λ = 0.001) and the middle range of symmetric division probability (symmetric division probabilities = 0.4 — 0.8), as can be seen in Fig 7.

**Figure 7.**
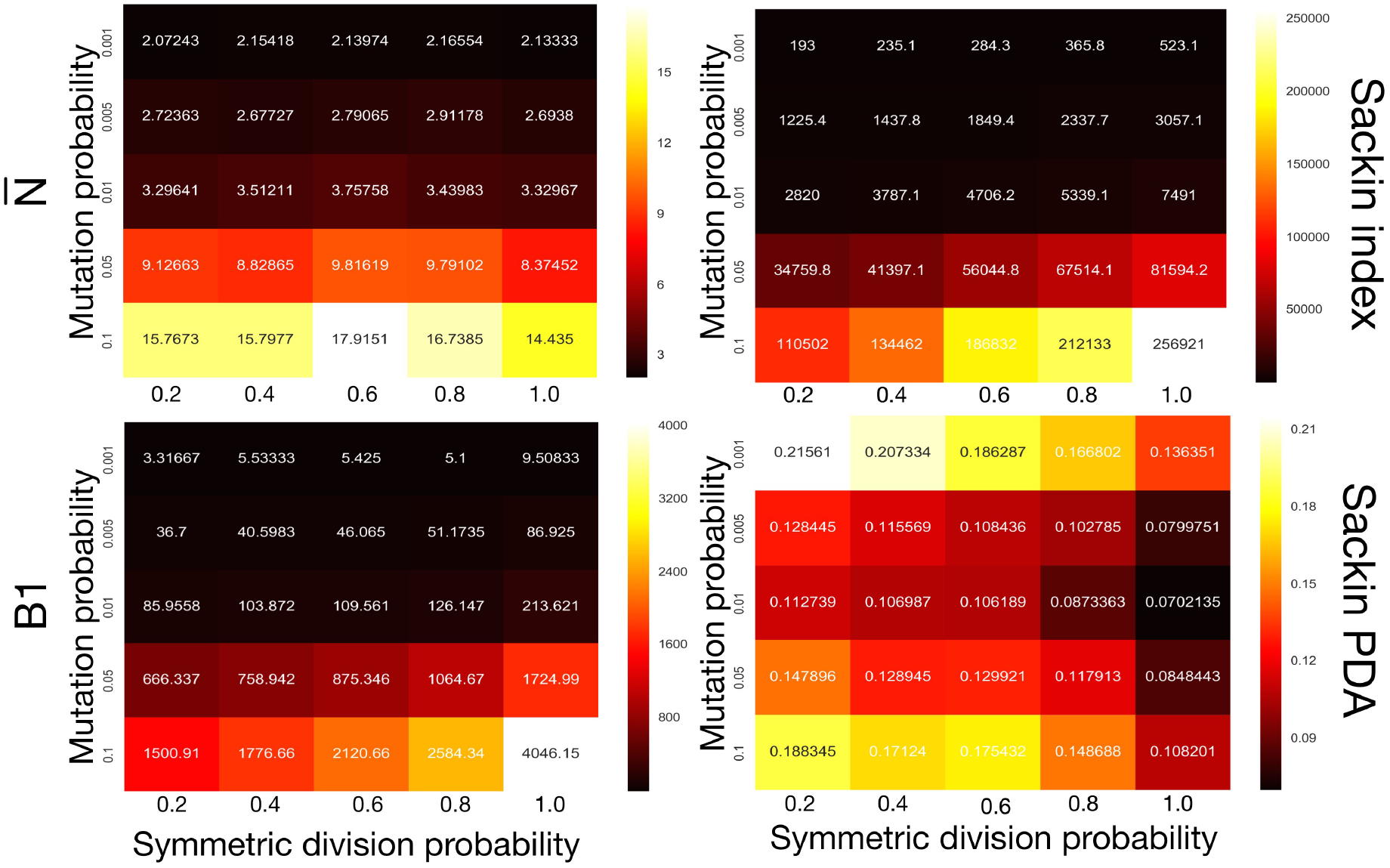
Comparing phylogenetic tree measures across symmetric division probability and mutation probability. We plot the average of each of four phylogenetic tree measures at the end of each of 10 simulations for a range of symmetric division probabilities and mutation probabilities. We vary mutational probability over two orders of magnitude (0.1 — 0.001), and simulate all tested symmetric division probabilities. Rank is maintained across symmetric division probabilities for each of the three of the four measures with which we could discriminate between symmetric division probabilities with changing mutation probability, allowing for differentiation between parameters. As before, the N statistic is not predictive. As expected, for the non-normalized indices, Sackin and B1, the measures change monotonically with both symmetric division and mutation probability. For the PDA normalized Sackin index, however, there is a global minimum for λ = 0.01 and *α* = 1.

## Discussion

While the use of phylogenetic trees is increasing in translational oncology laboratories, there has yet to be a method found by which we can utilise the information clinically. To address this shortcoming, we worked to leverage the growing interest in biomarker derivation from spatially distinct tumour biopsies^65^, and the recent success of Leventhal^39^ and others in teasing apart complex biological rules from phylogenetic information. We developed an individual based model of tumour growth under a TIC driven proliferative heterogeneity which undergoes neutral evolution. We then developed an algorithm to construct phylogenetic trees from simulated tumours. The resultant trees were then analysed and compared using a suite of statistical measures of tree (im)balance. Through this method, we have generated a large dataset that includes the observed statistical measures of the ‘true’ phylogeny for tumours with a range of symmetric division probabilities.

In particular, we compared the classical measures of tree topology – the Sackin index and the B1 statistic – as well as normalized versions of each across several parameters of our spatial and non-spatial models as well as through the process of tumour growth. Not surprisingly, we found that the Sackin index was able to discriminate between the families of simulations as it is directly correlated with branch number (in this case correlating with total number of mutations in the TICs, which also is increased with increasing symmetric division probability). Encouragingly, we also found that the normalised version of this metric was able to discriminate between the different symmetric division probabilities, suggesting a more meaningful (and measurable) topologic difference between the underlying phylogenetic trees resulting from these parameter changes (representing diverse biological traits).

While we have shown that these measures differ significantly from one another, we have not yet provided a method by which we can use the metric of a given tree to directly predict the symmetric division probability of an unknown tumour. However, the present work at least allows us to understand the rank order of symmetric division rate for two tumours given their measured indices. This could be particularly useful in certain clinical settings. For example, this could allow us to determine how a given therapy affects symmetric division probability by using our calculated measures over serial biopsies, and subsequent phylogenetic reconstruction.

## Conclusions

Aiming towards a translatable method by which to infer the symmetric division probability in solid tumours, we have identified several phylogenetic tree based measures that correlate with TIC symmetric division probability. We have found several measures which are able to discern differences in simulated tumours between symmetric division probabilities. These results are robust to changes in tumour size, specifically maintaining their rank throughout tumour growth. The rate of mutation does affect these results to some degree, but rank is maintained permitting comparison through time, or between tumours of similar size.

While there is some overlap amongst the measures when more than one parameter is varied, with information on mutation probability and tumour size, relative symmetric division probability can be estimated. we have only restricted our focus to measures of (im)balance, a basic property of phylogenetic trees based only on their branching topology. With more information, such as evolutionary branch lengths^66,67^ which are linked to the ‘speed’ of a tumour’s molecular clock^64^, some of these limitations could be obviated. Further, we have only considered neutral evolution. While most tumour evolution is likely neutral^48^, there is certainly evidence for non-neutrality in the form of driver and passenger mutations^47,68^, which would drastically affect the resulting phylogenetic trees^38^ – especially with intervening treatment regimens. How non-neutral evolution and treatment affect our measures remain avenues for future work.

## Acknowledgements

The authors thank Andrea Sottoriva, Trevor Graham and Helen Byrne for insightful comments and discussions. AGF is supported by a Vice-Chancellor’s Fellowship from the University of Sheffield.

## Supplemental Material

### Pruning trees does not affect rank of statistics

To visualize the trees more easily in Fig 3, we prune the leaves from each full tree. While this changes the absolute value of each of the tree-based measures, it does not affect their relative ranking. This suggests that each measure is capturing something fundamental about the biology as it appears invariant with tree size. This is corroborated by the results shown in Fig 6, indicating that the rank of each measure is stable over tumour growth.

### Effect size of symmetric division probability

To better understand the impact of the symmetric division probability on changes in results tree topology, rather than just use differences between families of simulations, we compute the regression slope, R^2^ and p-value of the regression line for each case. For the B1 statistic we find a regression slope of 142.64, R^2^ = 0.72, *p* = 1.74 × 10^−71^. For the Sackin index we find a regression slope of 5178.61, R^2^ = 0.871, *p* ≈ 0. For the Yule normalised Sackin index we find a regression slope of —2.380, R^2^ = 0.743, *p* = 3.25 × 10^−75^. For the 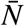 statistic we find a regression slope of –0.111, R^2^ = 0.0075, *p* = 0.172. These values are plotted in Fig 9.

**Figure 8.**
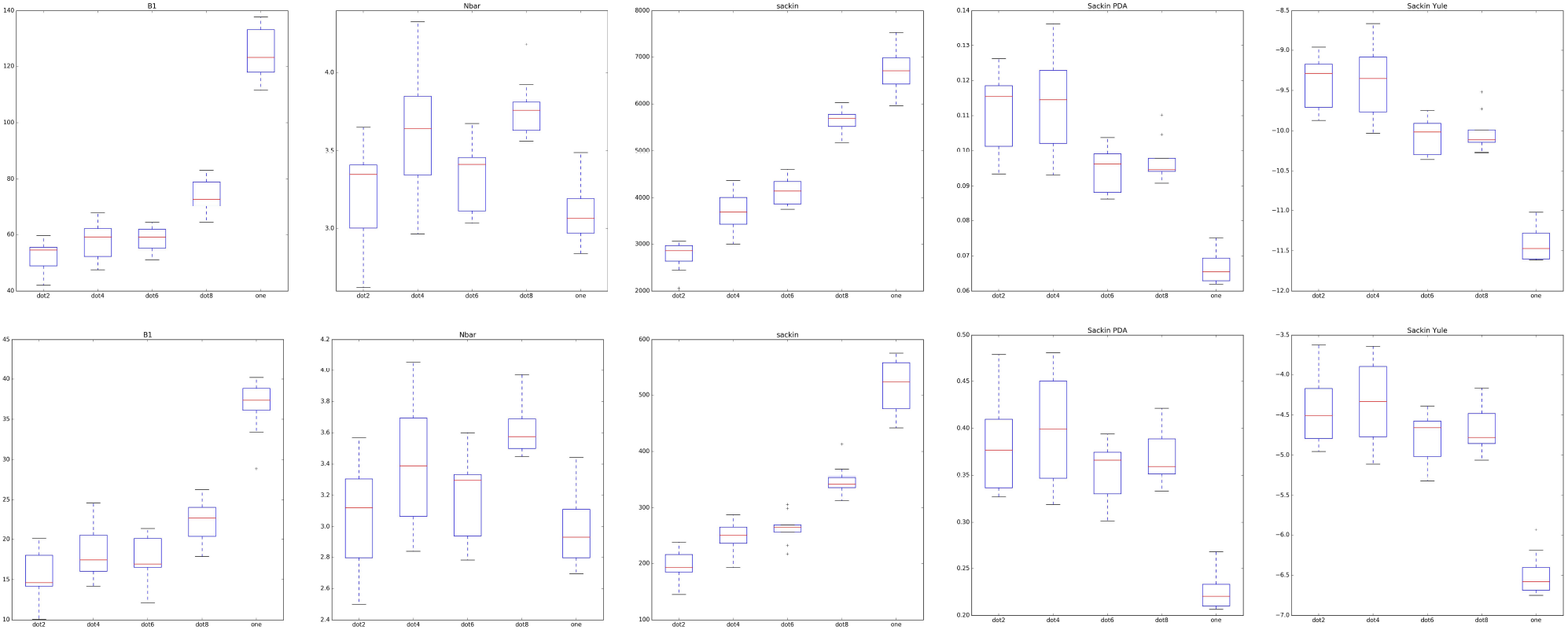
Raw and pruned trees give rise to qualitatively similar summary measures with rank preserved. For each tree-based measure considered in the main text, we plot the measure based on the full (upper) and pruned (lower) tree. For each pair, we plot the results from 10 simulations for each of the tested symmetric division probabilities. From left to right, we plot the B1 statistic, 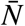, the Sackin index, the PDA normalised Sackin index and finally the Yule normalised Sackin index.

**Figure 9.**
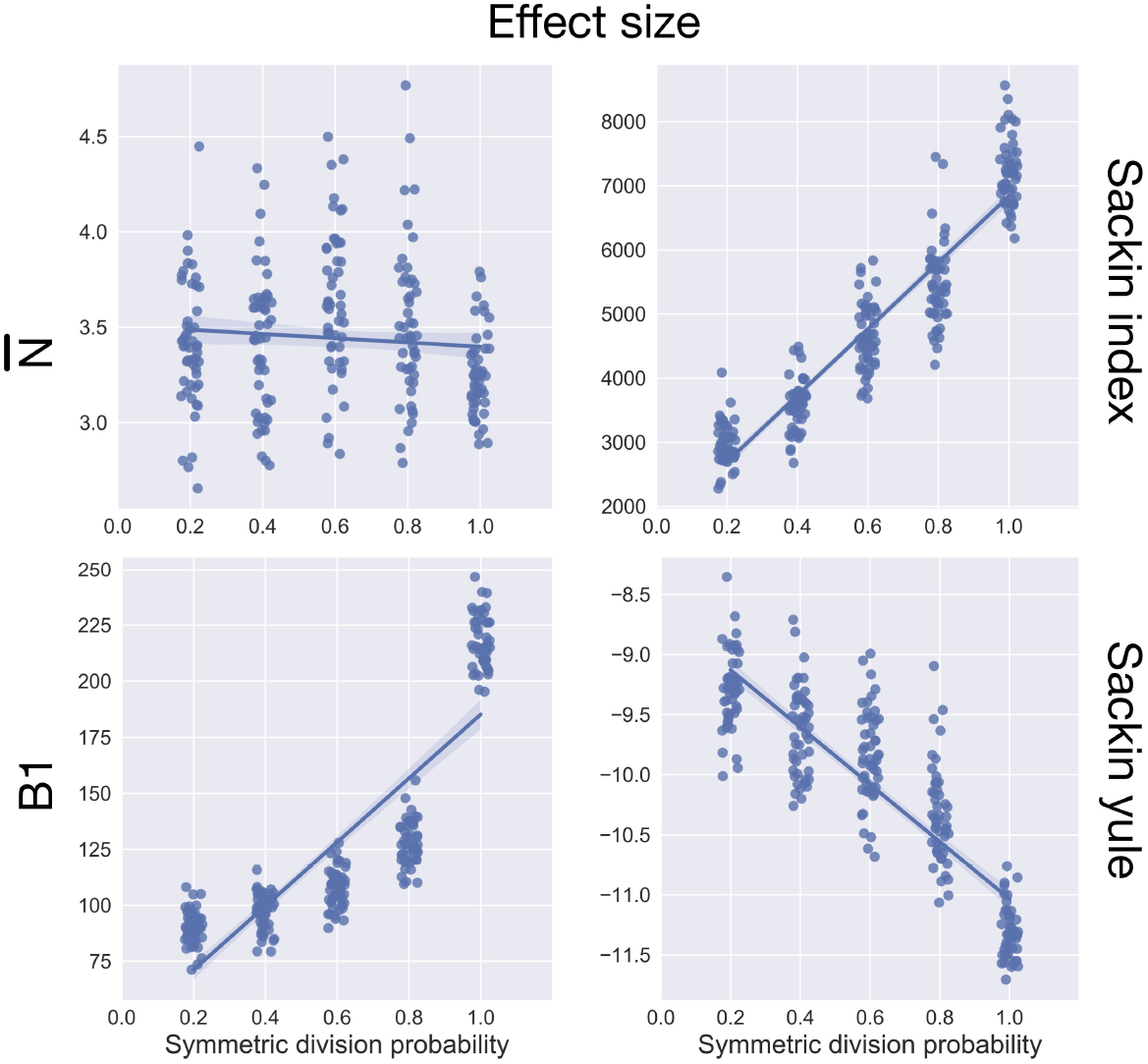
Effect size of symmetric division for four tree-based measures. We plot the effect size for the data shown in Fig 5.

### Algorithm for generating individual cell ‘genomes’ from mutational flag and life history

Here we describe the algorithm we created to develop the individual cells ‘genomes’ from the mutational flag and the life history. Using this reconstruction algorithm allows for significant increase in speed of our tumour growth model and reduced memory requirements by several orders of magnitude.

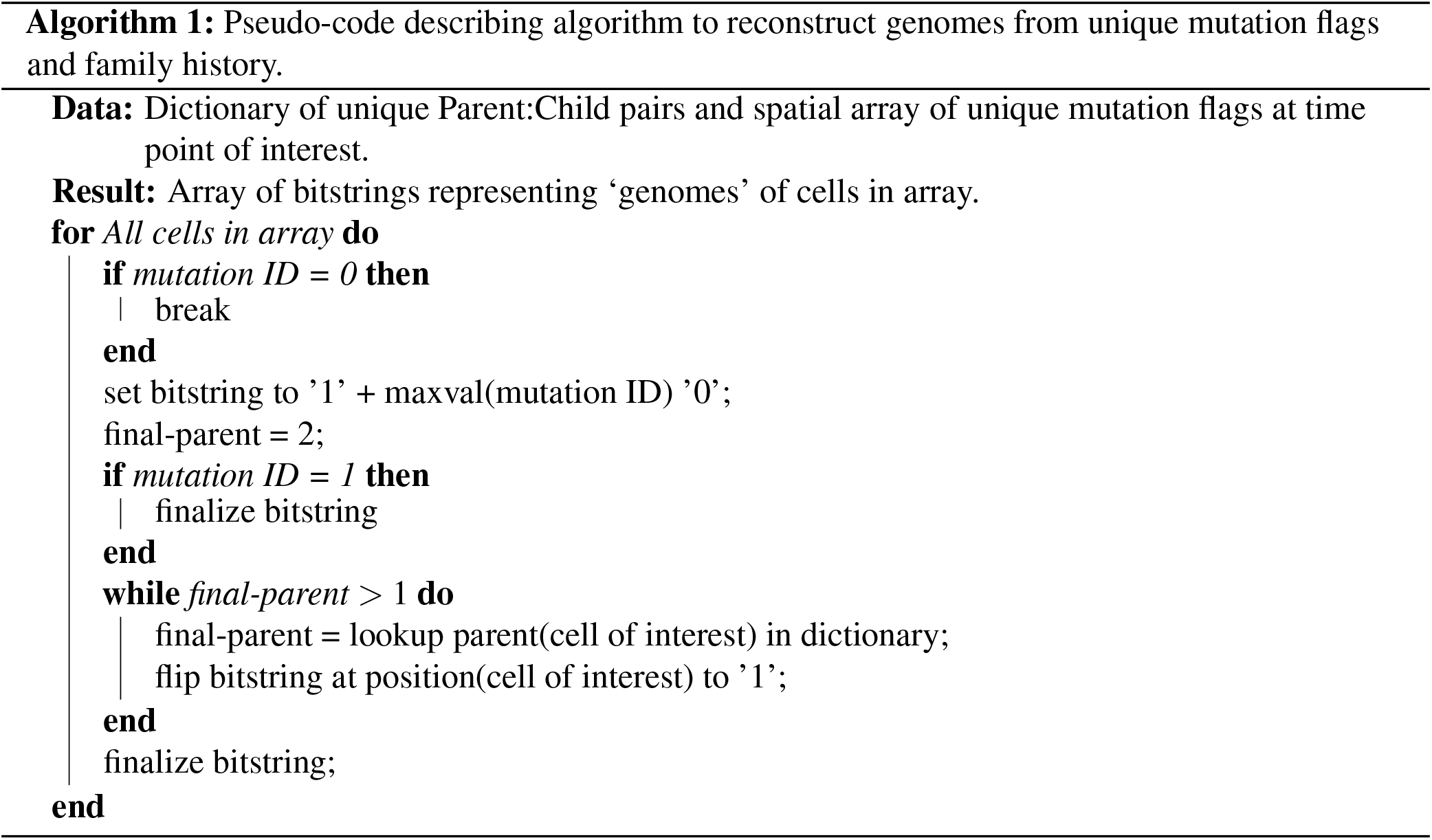

